## Supplementary Note and Table for "Surface protein imputation from single cell transcriptomes by deep neural networks"

**Supplementary note 1.** cTP-net models tested on CITE-seq CBMC data sets

Table S5 illustrates the different models we have tested. The first column indicates the differences to the finalized models, while the second column shows the correlation of the predicted protein abundance to the true protein abundance in the holdout setting on CITE-seq CBMC data set. As shown by Table S5, missing any component of the final model will result in inferior performance.

**Table S1.** Summary table of five data sets analyzed in this study

| Data | Technology | Cell population | # of subjects | # of cells | # of genes | # of proteins | # of cell types |
|---|---|---|---|---|---|---|---|
| **CITE-PBMC** | CITE-seq | PBMC | 1 | 7667 | 13517 | 10 | 8 |
| **CITE-CBMC** | CITE-seq | CBMC | 1 | 8005 | 14505 | 10 | 12 |
| **REAP-PBMC** | REAP-seq | PBMC | 1 | 4326 | 10811 | 10 | NA |
| **CITE-BMMC** | CITE-seq | BMMC | 1 | 33455 | 17009 | 25 | NA |
| **HCA-CBMC** | 10x | CBMC | 8 | 260,000 | 12611 | NA | NA |
| **HCA-BMMC** | 10x | BMMC | 8 | 270,000 | 12611 | NA | NA |

**Table S2.** Cell type summary of CITE-seq data sets

| Data | Cell types |
|---|---|
| **CITE-PBMC** | B, CD8 T-1, CD4 T, NK, DC, CD14+CD16+ Mono, CD14-CD16+ Mono, CD8 T 2 |
| **CITE-CBMC** | B, CD8 T, CD4 T, NK, DC, CD14+ Mono, CD16+ Mono, pDC, CD34+, Eryth, Unknown |

**Table S3.** Top 20 highest influence score genes for each protein in CITE-PBMC data set

| CD3 | CD4 | CD8 | CD2 | CD45RA | CD57 | CD16 | CD14 | CD11c | CD19 |
|---|---|---|---|---|---|---|---|---|---|
| *CD3D* | *CD8B* | *CD8B* | *CCL5* | *KLRB1* | *NUDT6* | *CHL1* | *C1orf115* | *CFD* | *CCL5* |
| *IL7R* | *CD8A* | *CD8A* | *IL7R* | *CCL5* | *MZT2A* | *RP11-242C19.2* | *PEAK1* | *MAL* | *CD8B* |
| *CD8B* | *RP11-291B21.2* | *CCL5* | *RP4-539M6.22* | *EIF1AX* | *ATP2A2* | *GCSH* | *ALDH7A1* | *ANKRD36C* | *RN7SL600P* |
| *FCER1G* | *CCL5* | *TRDC* | *LTV1* | *CD7* | *IQCE* | *NRL* | *CYBB* | *BLOC1S3* | *MYO1D* |
| *TRDC* | *NCR3* | *RP11-291B21.2* | *RP11-452L6.5* | *TST* | *PKNOX1* | *DBF4* | *ISYNA1* | *IGLL5* | *HSF2* |
| *AKR7A2* | *KLRB1* | *CHMP7* | *FBXO10* | *MFSD7* | *FBXW8* | *SPHK2* | *LMAN1* | *RP11-159G9.5* | *AC142528.1* |
| *HELLS* | *DDIT3* | *ZAP70* | *LINC00384* | *ZFAS1* | *CTA-217C2.1* | *CDKL1* | *FAM162A* | *ARMCX1* | *DNAJA3* |
| *ALG10* | *CTD-2547L16.1* | *BMP8B* | *ACAP2* | *TAPSAR1* | *CNOT11* | *TIMM21* | *SLC4A7* | *SLC6A16* | *GLB1L* |
| *FGD5-AS1* | *C18orf25* | *FAH* | *PPCDC* | *PLEKHF1* | *HSD17B4* | *MRPS18C* | *MIER3* | *LRRC16A* | *LIMD2* |
| *COMMD7* | *FPGT* | *AC009299.3* | *ANKRD39* | *CTBP1-AS2* | *CLEC4E* | *C7orf43* | *SLC11A2* | *TRAF1* | *DTX3L* |
| *CTA-292E10.8* | *NETO2* | *CMKLR1* | *AIM2* | *CYP27A1* | *FAM98C* | *GORASP2* | *ZAP70* | *PABPN1* | *PTCD2* |
| *ZC2HC1A* | *GDAP1* | *ENTPD1* | *TTLL12* | *MAN1A2* | *PRMT1* | *CTD-2555C10.3* | *MAP4* | *ADM* | *LPAR1* |
| *INADL* | *CSTF1* | *PIK3CA* | *GABBR1* | *FAM115C* | *SLC25A11* | *LEPROT* | *TTTY15* | *KIAA0319L* | *ZNF649* |
| *SHISA4* | *RP11-159H10.3* | *WDR7* | *DCUN1D4* | *CST3* | *TCEANC2* | *RUSC1* | *HS1BP3* | *MRPL4* | *HLA-DRB5* |
| *DCAF4* | *RP11-451M19.3* | *HEG1* | *CPD* | *NAIF1* | *LCTL* | *POLR2L* | *PRPSAP1* | *NDRG1* | *LIN54* |
| *HPGDS* | *ENTPD1-AS1* | *NPAT* | *RAPGEFL1* | *RP11-83N9.5* | *CAPN1* | *RP11-85A1.3* | *ZBTB38* | *FAM63A* | *USP32* |
| *PACSIN1* | *SLC4A10* | *7-Sep* | *U91328.20* | *FCGR3A* | *VPS26A* | *FKBP7* | *PIK3R1* | *RPL34* | *AIM2* |
| *ARID4B* | *FAAH2* | *CDT1* | *EIF4H* | *CCDC163P* | *ECHS1* | *RNF24* | *PIGG* | *FAM118B* | *SLC12A7* |
| *ATP11A* | *AP5B1* | *QRICH1* | *AC073115.7* | *POLR1C* | *FLVCR1-AS1* | *TBXAS1* | *DESI2* | *UBQLN4* | *ZNF671* |
| *RN7SL521P* | *DHPS* | *AP2M1* | *RP11-4O1.2* | *PRSS35* | *RP11-421L21.2* | *WDR83* | *SIRT5* | *FKBP15* | *TNNI2* |

**Table S4.** Gene set enrichment analysis on cell-immunophenotype pairs that cTP-net predict well in CITE-PBMC data set

| **Surface protein** | **Cell type** | **GO pathways** |
|---|---|---|
| **CD45RA** | **CD14-CD16+Mono** | GO_CATABOLIC_PROCESS |
| | | GO_PROTEIN_LOCALIZATION |
| | | GO_REGULATION_OF_CELLULAR_COMPONENT_BIOGENESIS |
| | | GO_CELLULAR_RESPONSE_TO_STRESS |
| | | GO_CELLULAR_RESPONSE_TO_DNA_DAMAGE_STIMULUS |
| | | GO_RNA_BINDING |
| | | GO_ESTABLISHMENT_OF_LOCALIZATION_IN_CELL |
| | | GO_CELL_CYCLE |
| | | GO_SINGLE_ORGANISM_BIOSYNTHETIC_PROCESS |
| | | GO_CELLULAR_MACROMOLECULE_LOCALIZATION |
| **CD11c** | **CD14-CD16+Mono** | GO_CELLULAR_RESPONSE_TO_STRESS |
| | | GO_NEGATIVE_REGULATION_OF_GENE_EXPRESSION |
| | | GO_POSITIVE_REGULATION_OF_BIOSYNTHETIC_PROCESS |
| | | GO_POSITIVE_REGULATION_OF_GENE_EXPRESSION |
| | | GO_CELL_CYCLE |
| | | GO_POSITIVE_REGULATION_OF_PROTEIN_METABOLIC_PROCESS |
| | | GO_NEGATIVE_REGULATION_OF_NITROGEN_COMPOUND_METABOLIC_PROCESS |
| | | GO_CYTOSKELETON |
| | | GO_CHROMOSOME |
| | | GO_ENZYME_BINDING |
| **CD45RA** | **CD8 T 2** | GO_ENZYME_BINDING |
| | | GO_RNA_BINDING |
| | | GO_RIBONUCLEOPROTEIN_COMPLEX |
| | | GO_REGULATION_OF_TRANSCRIPTION_FROM_RNA_POLYMERASE_II_PROMOTER |
| | | GO_CELL_CYCLE |
| | | GO_RNA_PROCESSING |
| | | GO_POSITIVE_REGULATION_OF_BIOSYNTHETIC_PROCESS |
| | | GO_CYTOSKELETON |
| | | GO_RIBONUCLEOTIDE_BINDING |
| | | GO_POSITIVE_REGULATION_OF_GENE_EXPRESSION |
| **CD45RA** | **CD4 T** | GO_REGULATION_OF_IMMUNE_SYSTEM_PROCESS |
| | | GO_IMMUNE_SYSTEM_PROCESS |
| | | GO_VACUOLE |
| | | GO_SMALL_MOLECULE_METABOLIC_PROCESS |
| | | GO_ORGANONITROGEN_COMPOUND_METABOLIC_PROCESS |
| | | GO_ESTABLISHMENT_OF_LOCALIZATION_IN_CELL |
| | | GO_POSITIVE_REGULATION_OF_MULTICELLULAR_ORGANISMAL_PROCESS |

| | | |
|---|---|---|
| | | GO_ENDOPLASMIC_RETICULUM |
| | | GO_REGULATION_OF_TRANSCRIPTION_FROM_RNA_POLYMERASE_II_PROMOTER |
| | | GO_PROTEIN_LOCALIZATION |
| **CD11c** | **CD14+CD16 + Mono** | GO_POSITIVE_REGULATION_OF_GENE_EXPRESSION |
| | | GO_DNA_REPLICATION |
| | | GO_POSITIVE_REGULATION_OF_MOLECULAR_FUNCTION |
| | | GO_POSITIVE_REGULATION_OF_BIOSYNTHETIC_PROCESS |
| | | GO_SINGLE_ORGANISM_BIOSYNTHETIC_PROCESS |
| | | GO_DNA_DEPENDENT_DNA_REPLICATION |
| | | GO_PHOSPHATE_CONTAINING_COMPOUND_METABOLIC_PROCESS |
| | | GO_CELL_JUNCTION |
| | | GO_CYTOKINE_RECEPTOR_BINDING |
| | | GO_ORGANONITROGEN_COMPOUND_BIOSYNTHETIC_PROCESS |
| **CD45RA** | **DC** | GO_NEGATIVE_REGULATION_OF_NITROGEN_COMPOUND_METABOLIC_PROCESS |
| | | GO_POLY_A_RNA_BINDING |
| | | GO_CHROMOSOME_ORGANIZATION |
| | | GO_REGULATION_OF_DNA_METABOLIC_PROCESS |
| | | GO_RNA_BINDING |
| | | GO_MACROMOLECULAR_COMPLEX_BINDING |
| | | GO_PHOSPHATE_CONTAINING_COMPOUND_METABOLIC_PROCESS |
| | | GO_NEGATIVE_REGULATION_OF_GENE_EXPRESSION |
| | | GO_ESTABLISHMENT_OF_LOCALIZATION_IN_CELL |
| | | GO_DNA_METABOLIC_PROCESS |
| **CD11c** | **DC** | GO_ENZYME_BINDING |
| | | GO_RIBONUCLEOTIDE_BINDING |
| | | GO_ESTABLISHMENT_OF_LOCALIZATION_IN_CELL |
| | | GO_NEGATIVE_REGULATION_OF_PROTEIN_METABOLIC_PROCESS |
| | | GO_IMMUNE_SYSTEM_PROCESS |
| | | GO_ORGANONITROGEN_COMPOUND_BIOSYNTHETIC_PROCESS |
| | | GO_PHOSPHATE_CONTAINING_COMPOUND_METABOLIC_PROCESS |
| | | GO_PHOSPHORYLATION |
| | | GO_NEGATIVE_REGULATION_OF_PROTEIN_MODIFICATION_PROCESS |
| | | GO_TRANSFERASE_ACTIVITY_TRANSFERRING_PHOSPHORUS_CONTAINING_GROUPS |

**Table S5.** Summary table of different cTP-net models

| **Differences to the finalized model** | **Correlation** |
|---|---|
| Without SAVER-X denoising, without MB structure | 0.961±0.0004 |
| Without MB structure | 0.968±0.0005 |
| Without SAVER-X denoising | 0.959 ±0.0005 |
| L2 loss | 0.969±0.0002 |
| Set bottle neck layer to 256 nodes (128 in final model) | 0.968±0.0003 |
| Set bottle neck layer to 64 nodes (128 in final model) | 0.968±0.0003 |
| With additional shared layers | 0.969±0.0004 |
| With SeLU activation function | 0.966±0.0002 |
| With Dropout layer between layer1 and layer2 | 0.966±0.001 |
| **Final model** | **0.970**±0.0003 |
