## Supplementary Figure for "Surface protein imputation from single cell transcriptomes by deep neural networks"

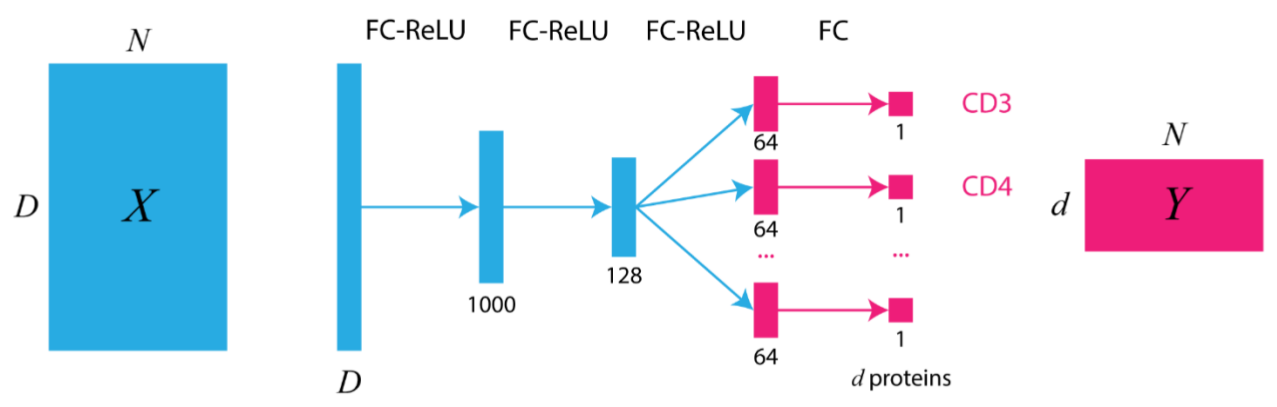


| |
|---|
| Supplementary Figure 1 |
| Neural network architecture of the cTP-net. |

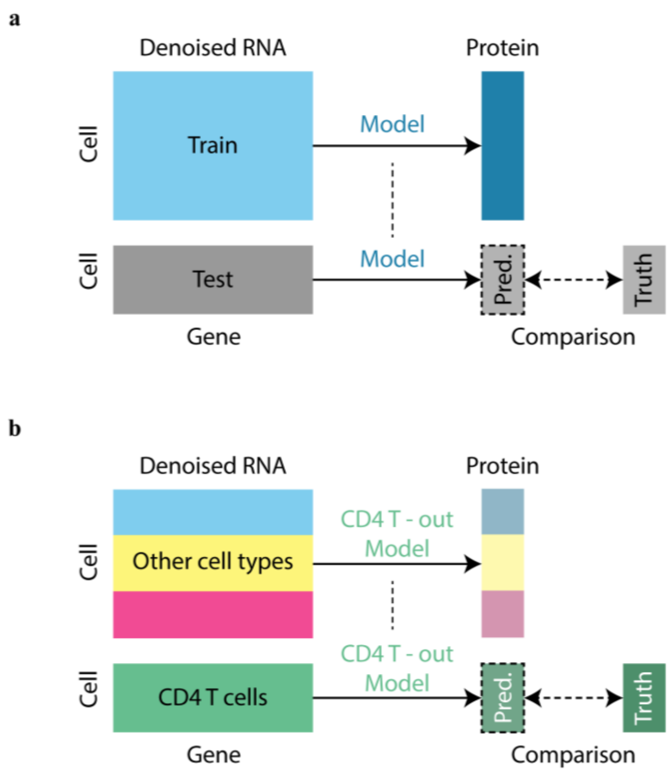


Supplementary Figure 2

Benchmark procedure.

(a) Holdout method validation scheme. (b) Out-of-cell-type benchmark scheme.

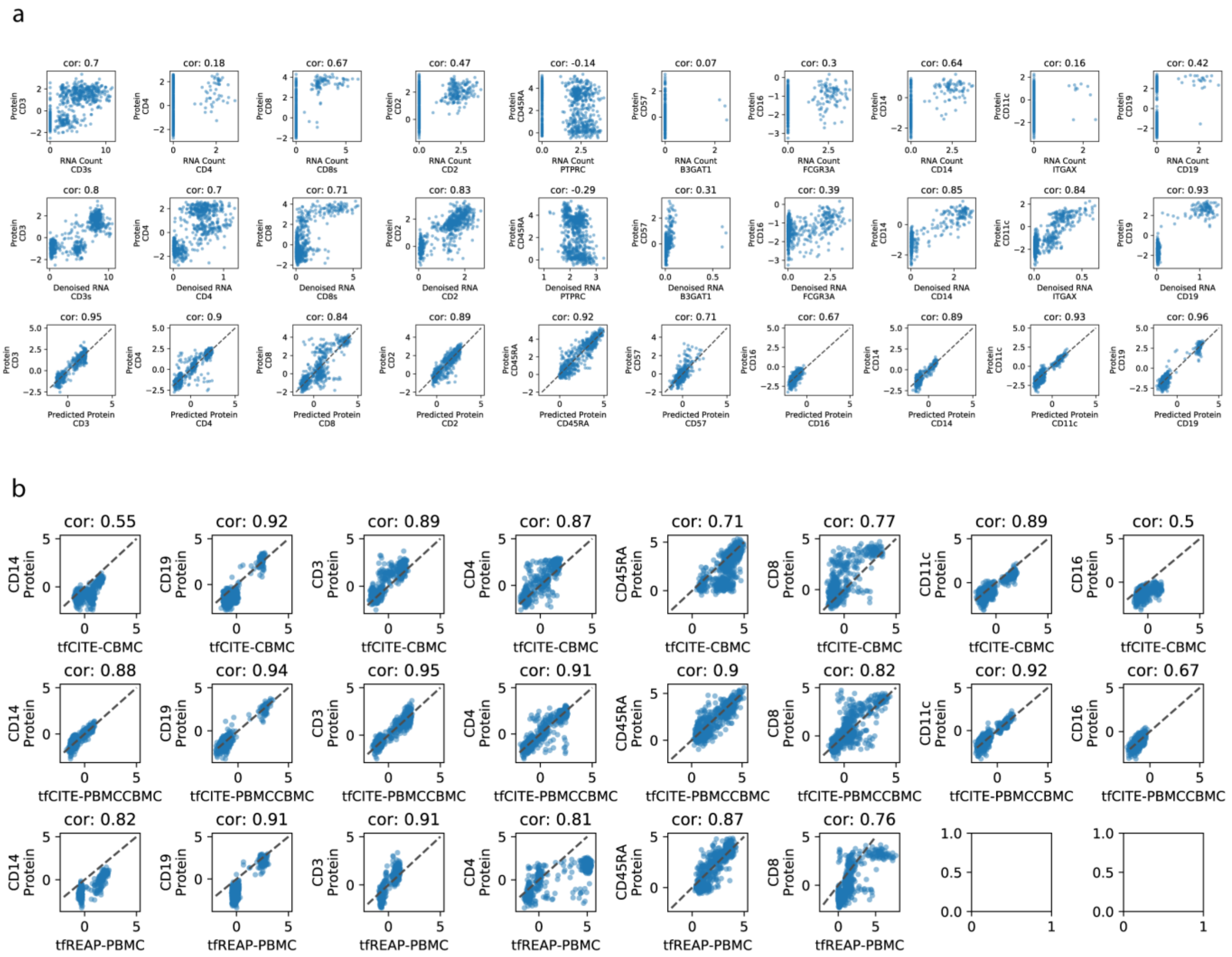
a
cor: 0.7
cor: 0.18
cor: 0.67
cor: 0.47
cor: -0.14
cor: 0.07
cor: 0.3
cor: 0.64
cor: 0.16
cor: 0.42
Protein CD3
Protein CD4
Protein CD8
Protein CD2
Protein CD45RA
Protein CD57
Protein CD16
Protein CD14
Protein CD11c
Protein CD19
RNA Count CD3s
RNA Count CD4
RNA Count CD8s
RNA Count CD2
RNA Count PTPRC
RNA Count B3GAT1
RNA Count FCGR3A
RNA Count CD14
RNA Count ITGAX
RNA Count CD19
cor: 0.8
cor: 0.7
cor: 0.71
cor: 0.83
cor: -0.29
cor: 0.31
cor: 0.39
cor: 0.85
cor: 0.84
cor: 0.93
Denoised RNA CD3s
Denoised RNA CD4
Denoised RNA CD8s
Denoised RNA CD2
Denoised RNA PTPRC
Denoised RNA B3GAT1
Denoised RNA FCGR3A
Denoised RNA CD14
Denoised RNA ITGAX
Denoised RNA CD19
cor: 0.95
cor: 0.9
cor: 0.84
cor: 0.89
cor: 0.92
cor: 0.71
cor: 0.67
cor: 0.89
cor: 0.93
cor: 0.96
Predicted Protein CD3
Predicted Protein CD4
Predicted Protein CD8
Predicted Protein CD2
Predicted Protein CD45RA
Predicted Protein CD57
Predicted Protein CD16
Predicted Protein CD14
Predicted Protein CD11c
Predicted Protein CD19
b
cor: 0.55
cor: 0.92
cor: 0.89
cor: 0.87
cor: 0.71
cor: 0.77
cor: 0.89
cor: 0.5
CD14 Protein
CD19 Protein
CD3 Protein
CD4 Protein
CD45RA Protein
CD8 Protein
CD11c Protein
CD16 Protein
tfCITE-CBMC
cor: 0.88
cor: 0.94
cor: 0.95
cor: 0.91
cor: 0.9
cor: 0.82
cor: 0.92
cor: 0.67
tfCITE-PBMCCBMC
cor: 0.82
cor: 0.91
cor: 0.91
cor: 0.81
cor: 0.87
cor: 0.76
tfREAP-PBMC

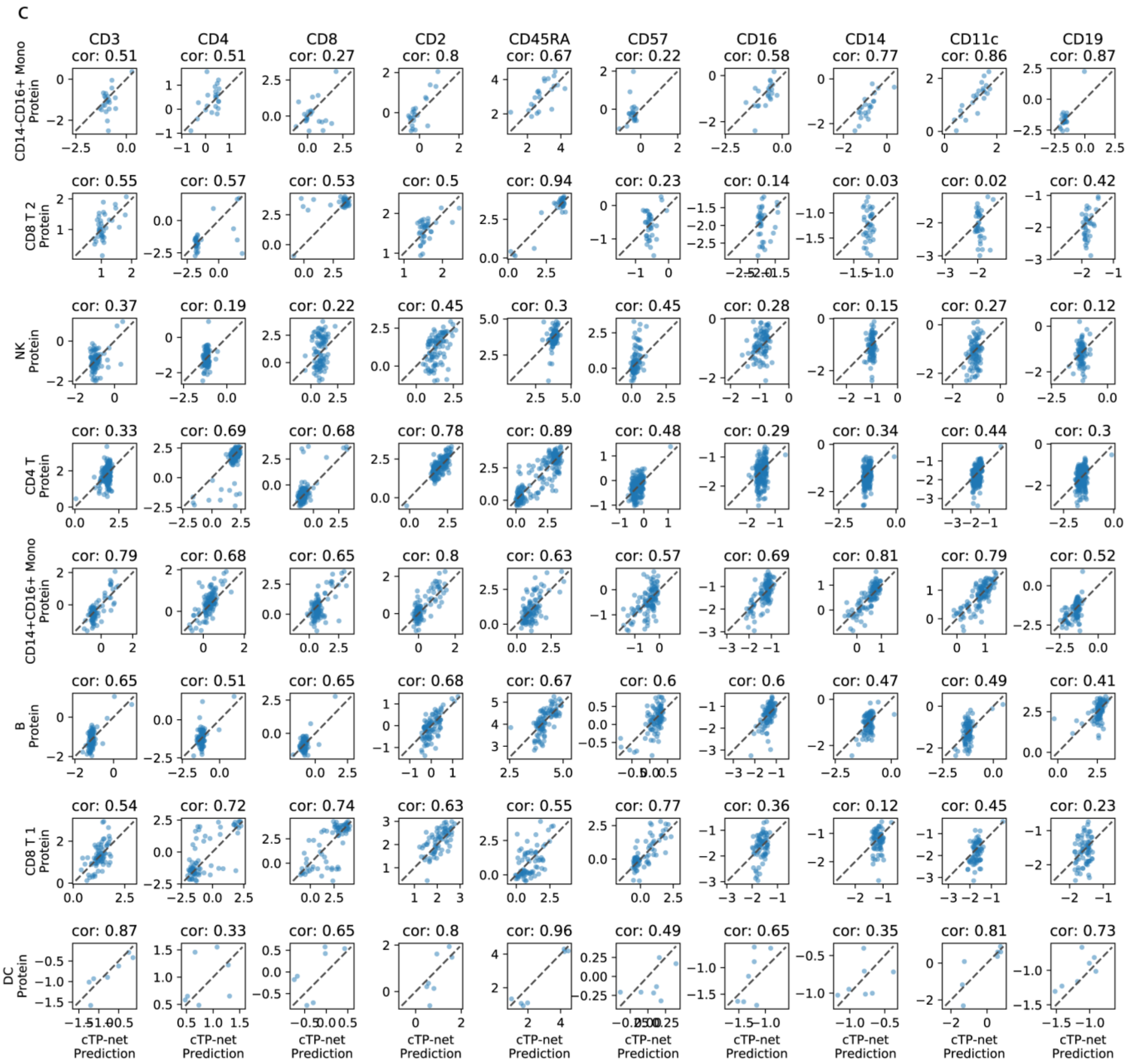


Supplementary Figure 3

Benchmark evaluation of cTP-net on CITE-PBMC data set.

(a) Benchmark correlation of true protein level vs. (1) Raw RNA count, (2) SAVER-X denoised RNA level, and (3) cTP-net predicted protein abundance in holdout method. (b) Benchmark correlation of truth protein level vs. (1) transfer learning from CITE-CBMC, (2) transfer learning from CITE-PBMCCBMC, and (3) transfer learning from REAP-PBMC. (c) Benchmark correlation of true protein level vs. cTP-net prediction in holdout method for each cell type.

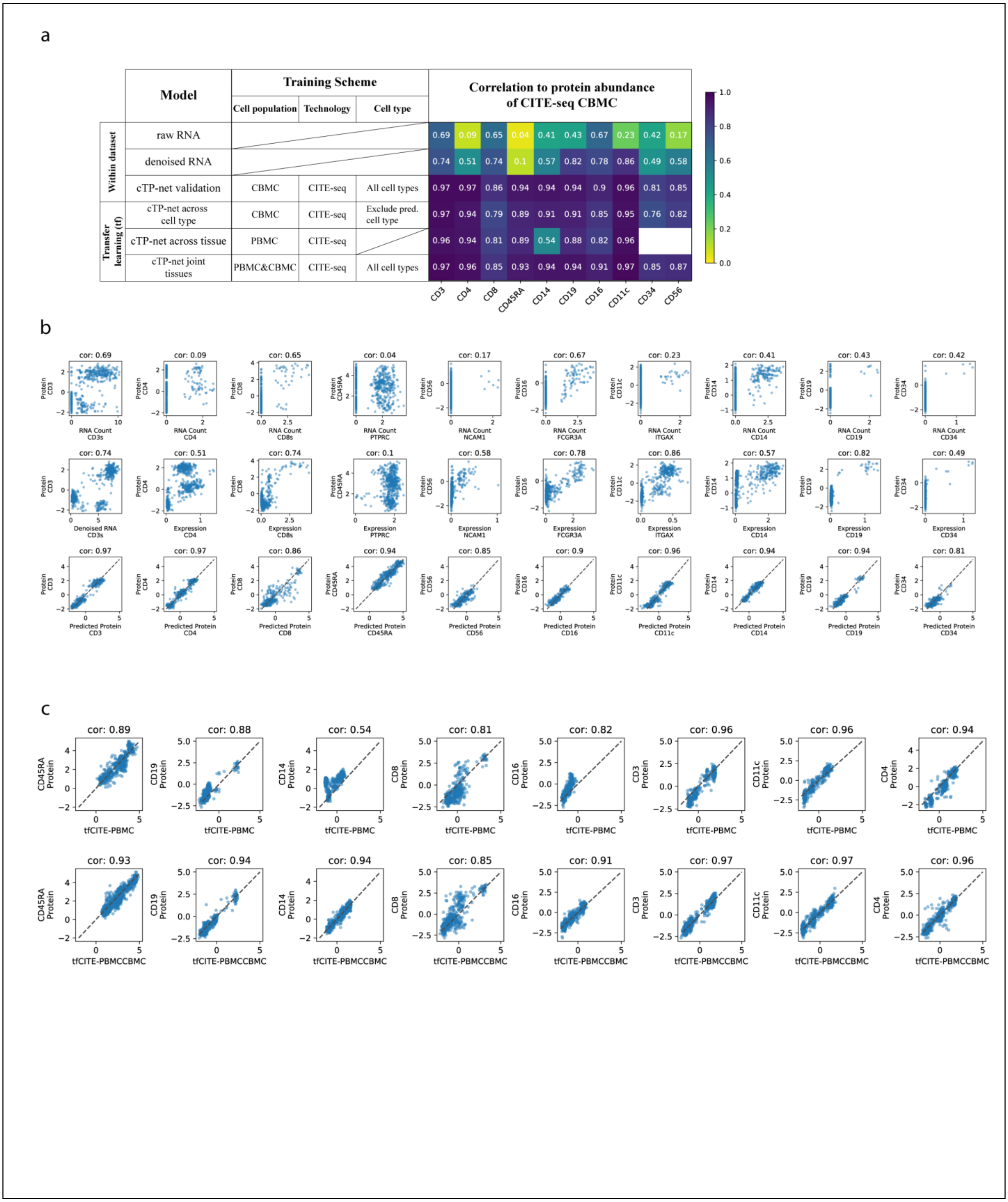
a
Model
Training Scheme
Cell population
Technology
Cell type
Correlation to protein abundance of CITE-seq CBMC
Within dataset
Transfer learning (tf)
raw RNA 0.69 0.09 0.65 0.04 0.41 0.43 0.67 0.23 0.42 0.17
denoised RNA 0.74 0.51 0.74 0.1 0.57 0.82 0.78 0.86 0.49 0.58
cTP-net validation CBMC CITE-seq All cell types 0.97 0.97 0.86 0.94 0.94 0.94 0.9 0.96 0.81 0.85
cTP-net across cell type CBMC CITE-seq Exclude pred. cell type 0.97 0.94 0.79 0.89 0.91 0.91 0.85 0.95 0.76 0.82
cTP-net across tissue PBMC CITE-seq 0.96 0.94 0.81 0.89 0.54 0.88 0.82 0.96
cTP-net joint tissues PBMC&CBMC CITE-seq All cell types 0.97 0.96 0.85 0.93 0.94 0.94 0.91 0.97 0.85 0.87
CD3 CD4 CD8 CD45RA CD14 CD19 CD16 CD11c CD34 CD56
b
cor: 0.69 cor: 0.09 cor: 0.65 cor: 0.04 cor: 0.17 cor: 0.67 cor: 0.23 cor: 0.41 cor: 0.43 cor: 0.42
RNA Count CD3s, CD4, CD8s, PTPRC, NCAM1, FCGR3A, ITGAX, CD14, CD19, CD34
cor: 0.74 cor: 0.51 cor: 0.74 cor: 0.1 cor: 0.58 cor: 0.78 cor: 0.86 cor: 0.57 cor: 0.82 cor: 0.49
Denoised RNA CD3s
Expression CD4, CD8s, PTPRC, NCAM1, FCGR3A, ITGAX, CD14, CD19, CD34
cor: 0.97 cor: 0.97 cor: 0.86 cor: 0.94 cor: 0.85 cor: 0.9 cor: 0.96 cor: 0.94 cor: 0.94 cor: 0.81
Predicted Protein CD3, CD4, CD8, CD45RA, CD56, CD16, CD11c, CD14, CD19, CD34
Protein CD3, CD4, CD8, CD45RA, CD56, CD16, CD11c, CD14, CD19, CD34
c
cor: 0.89 cor: 0.88 cor: 0.54 cor: 0.81 cor: 0.82 cor: 0.96 cor: 0.96 cor: 0.94
CD45RA Protein, CD19 Protein, CD14 Protein, CD8 Protein, CD16 Protein, CD3 Protein, CD11c Protein, CD4 Protein
tfCITE-PBMC
cor: 0.93 cor: 0.94 cor: 0.94 cor: 0.85 cor: 0.91 cor: 0.97 cor: 0.97 cor: 0.96
tfCITE-PBMCCBMC

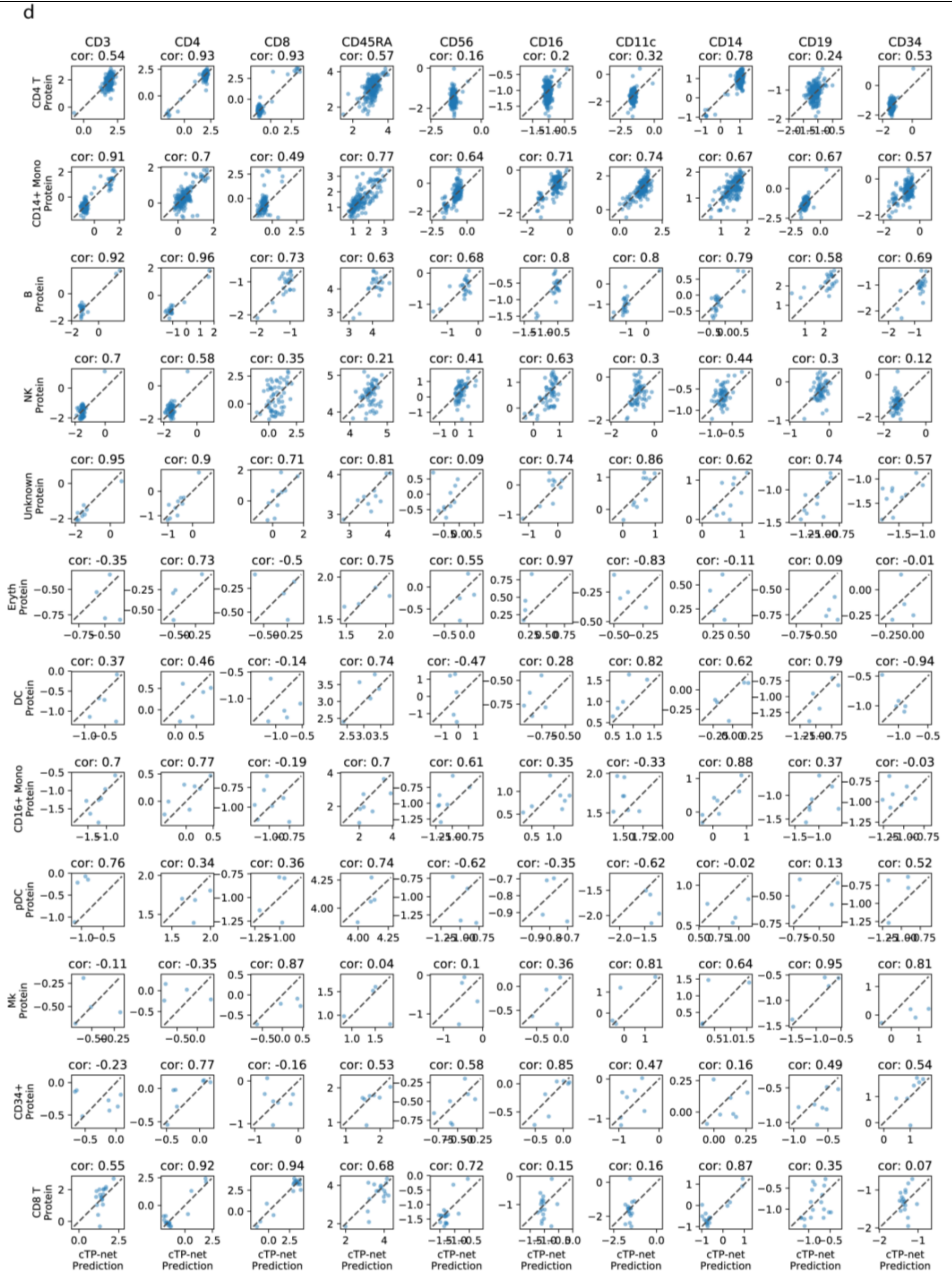


Supplementary Figure 4

Benchmark evaluation of cTP-net on CITE-CBMC data set.

**(a)** Benchmark evaluation heatmap of cTP-net and comparison with Seurat v3. The table on the left captures the detailed training scheme and model name of each test. **(b)** Benchmark correlation of true protein level vs. (1) Raw RNA count, (2) SAVER-X denoised RNA level, and (3) cTP-net predicted protein abundance in holdout method. **(c)** Benchmark correlation of truth protein level vs. (1) transfer learning from CITE-PBMC, and (2) transfer learning from CITE-PBMCCBMC. **(d)** Benchmark correlation of true protein level vs. cTP-net prediction in holdout method for each cell type.

**a**

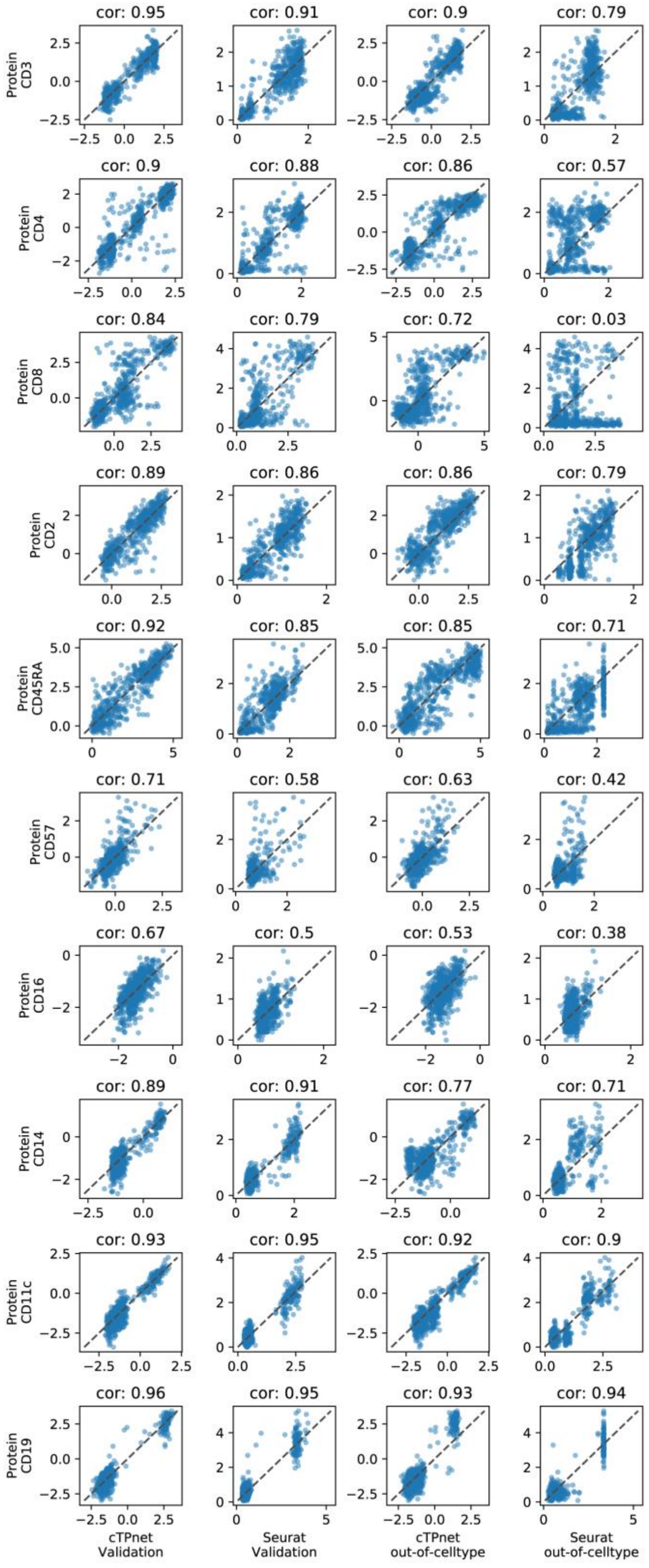
cor: 0.95
cor: 0.91
cor: 0.9
cor: 0.79
Protein CD3
cor: 0.9
cor: 0.88
cor: 0.86
cor: 0.57
Protein CD4
cor: 0.84
cor: 0.79
cor: 0.72
cor: 0.03
Protein CD8
cor: 0.89
cor: 0.86
cor: 0.86
cor: 0.79
Protein CD2
cor: 0.92
cor: 0.85
cor: 0.85
cor: 0.71
Protein CD45RA
cor: 0.71
cor: 0.58
cor: 0.63
cor: 0.42
Protein CD57
cor: 0.67
cor: 0.5
cor: 0.53
cor: 0.38
Protein CD16
cor: 0.89
cor: 0.91
cor: 0.77
cor: 0.71
Protein CD14
cor: 0.93
cor: 0.95
cor: 0.92
cor: 0.9
Protein CD11c
cor: 0.96
cor: 0.95
cor: 0.93
cor: 0.94
Protein CD19
cTPnet Validation
Seurat Validation
cTPnet out-of-celltype
Seurat out-of-celltype

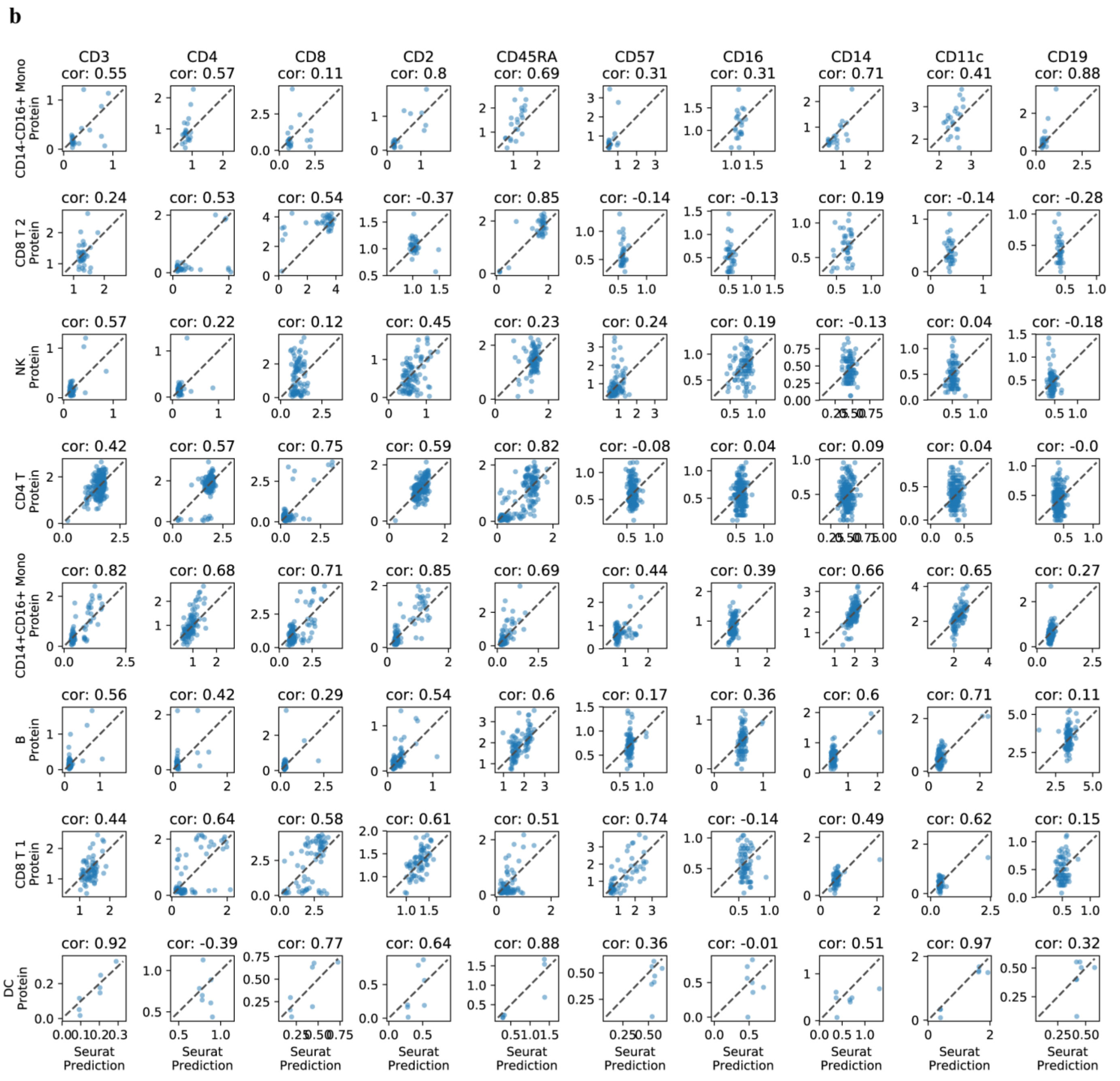


Supplementary Figure 5

Benchmark evaluation of Seurat v3 on CITE-PBMC data set.

**(a)** Benchmark correlation of true protein level vs. (1) cTP-net predicted protein abundance in holdout method, (2) Seurat v3 predicted protein abundance in holdout method, (3) out-of-cell-type cTP-net predicted protein abundance, and (4) out-of-cell-type Seurat v3 predicted protein abundance. **(b)** Benchmark correlation of truth protein level vs. (1) transfer learning from CITE-PBMC, and (2) transfer learning from CITE-PBMCCBMC. **(c)** Benchmark correlation of true protein level vs. cTP-net prediction in holdout method for each cell type.

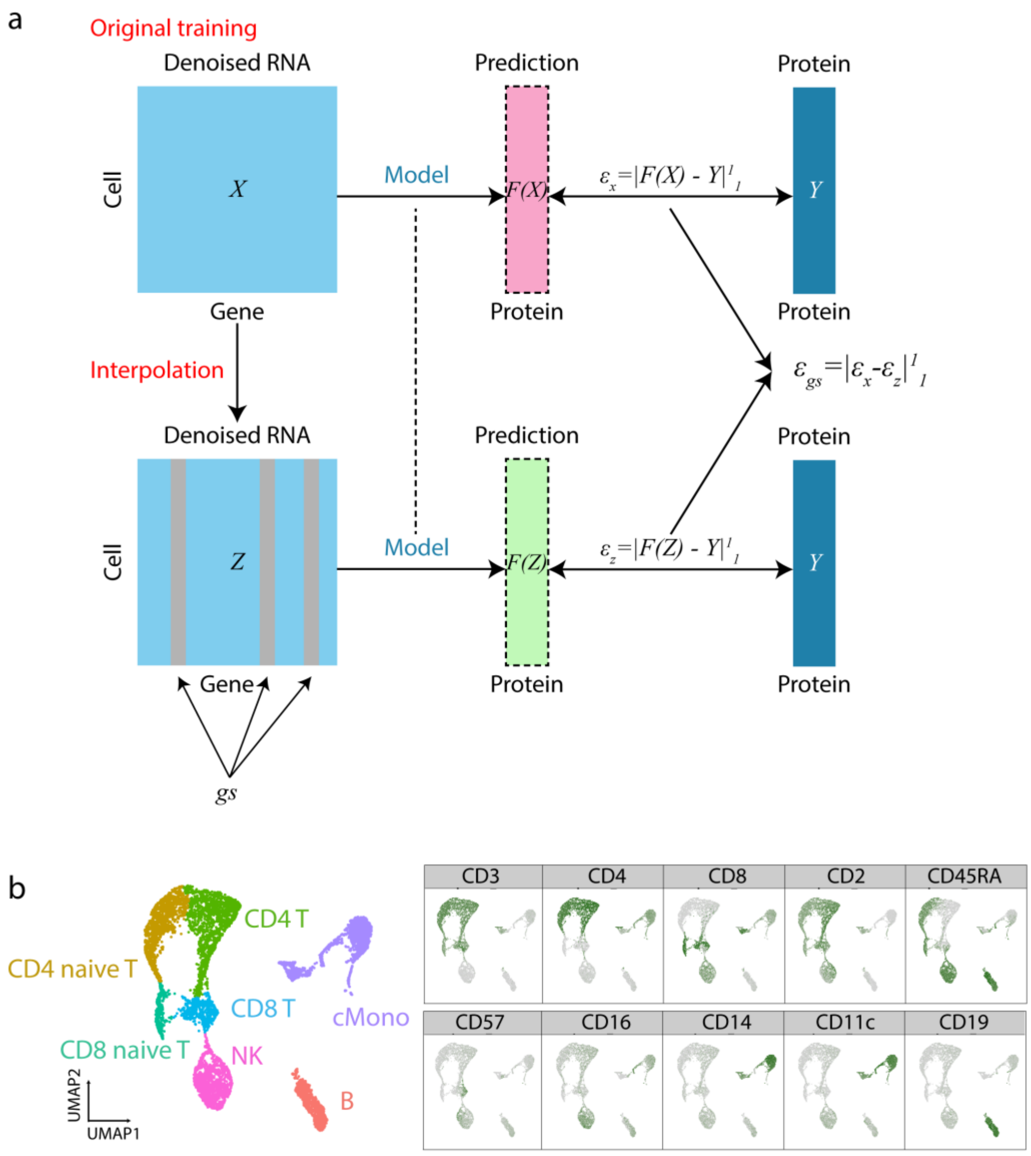


**Supplementary Figure 6.**

| |
|---|
| Supplementary Figure 6 |
| Interpolation analysis. |
| **(a)** Interpolation procedure in identify permutation based importance score for each gene in each protein prediction. **(b)** Dimension reduction analysis on the bottleneck layer on cTP-net trained on PBMCs from CITE-seq. |

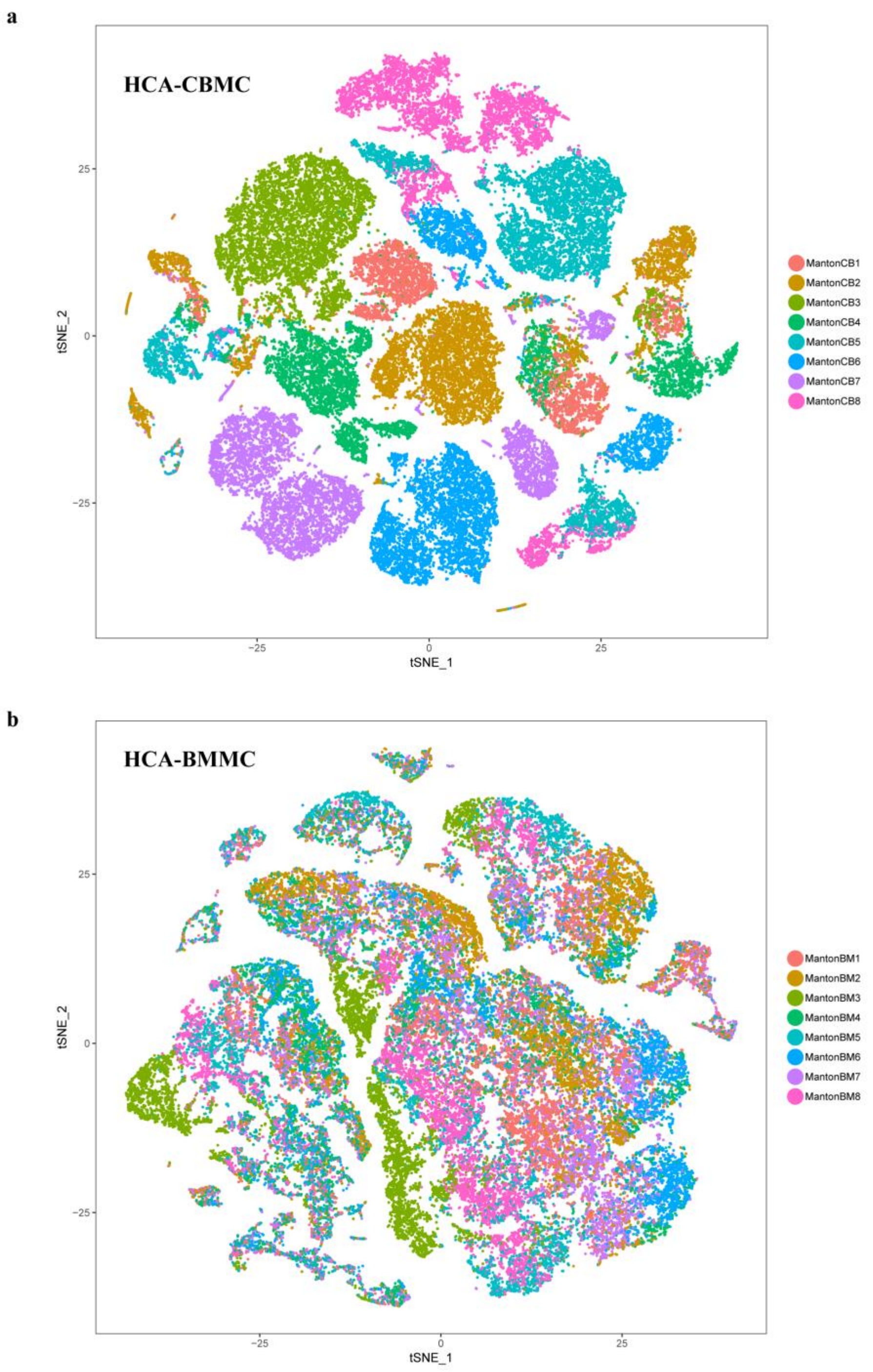


Supplementary Figure 7

Human Cell Atlas t-SNE plot based on normalized expression.

**(a)** t-SNE plot on Human Cell Atlas CBMCs based on normalized expression. Color indicates sample IDs. **(b)** t-SNE plot on Human Cell Atlas BMMCs based on normalized expression. Color indicates sample IDs. Strong batch effects observed in both data sets.

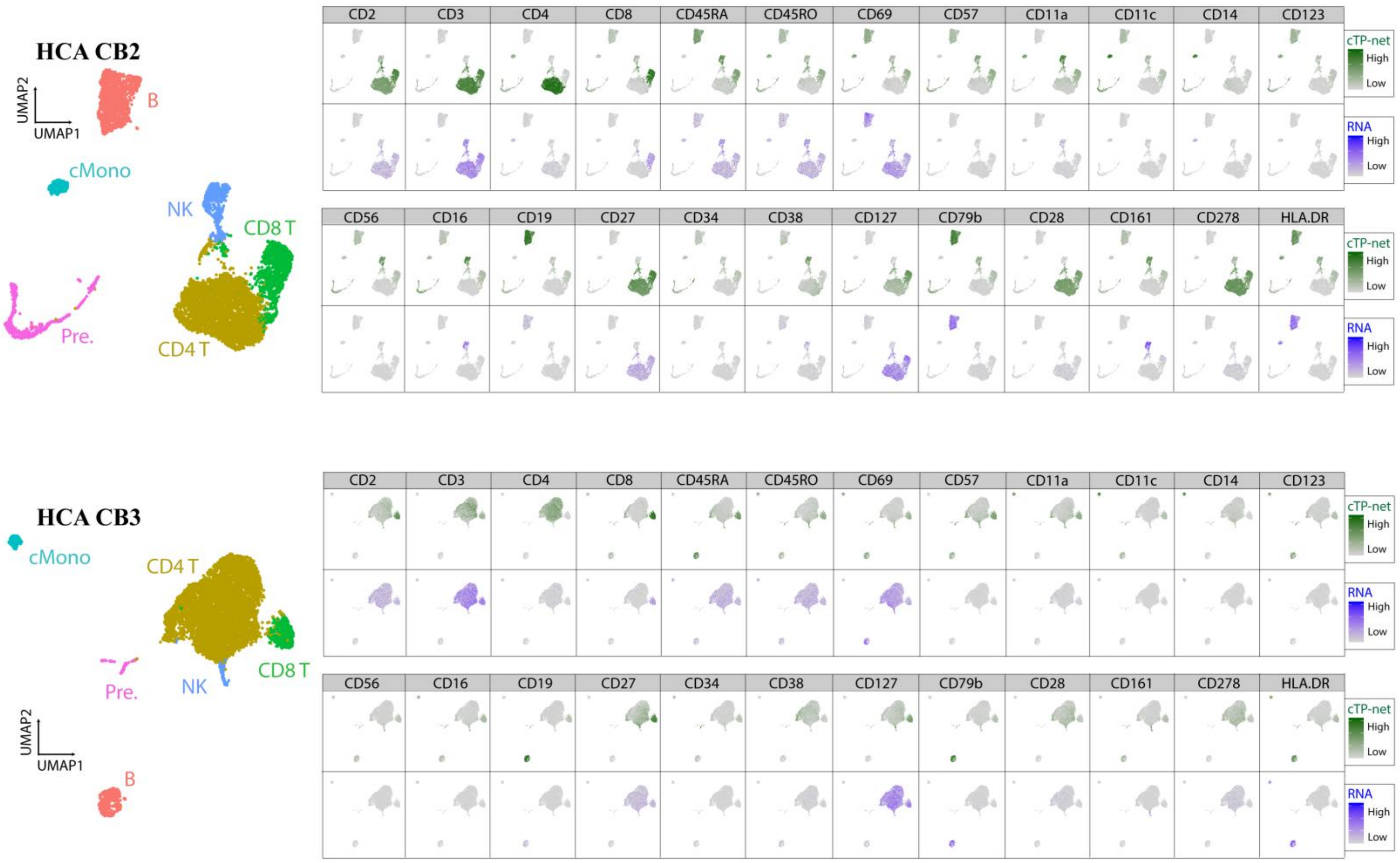

HCA CB2
UMAP2
UMAP1
B
cMono
NK
CD8 T
CD4 T
Pre.
CD2
CD3
CD4
CD8
CD45RA
CD45RO
CD69
CD57
CD11a
CD11c
CD14
CD123
cTP-net
High
Low
RNA
High
Low
CD56
CD16
CD19
CD27
CD34
CD38
CD127
CD79b
CD28
CD161
CD278
HLA.DR
HCA CB3
cMono
CD4 T
CD8 T
NK
Pre.
B

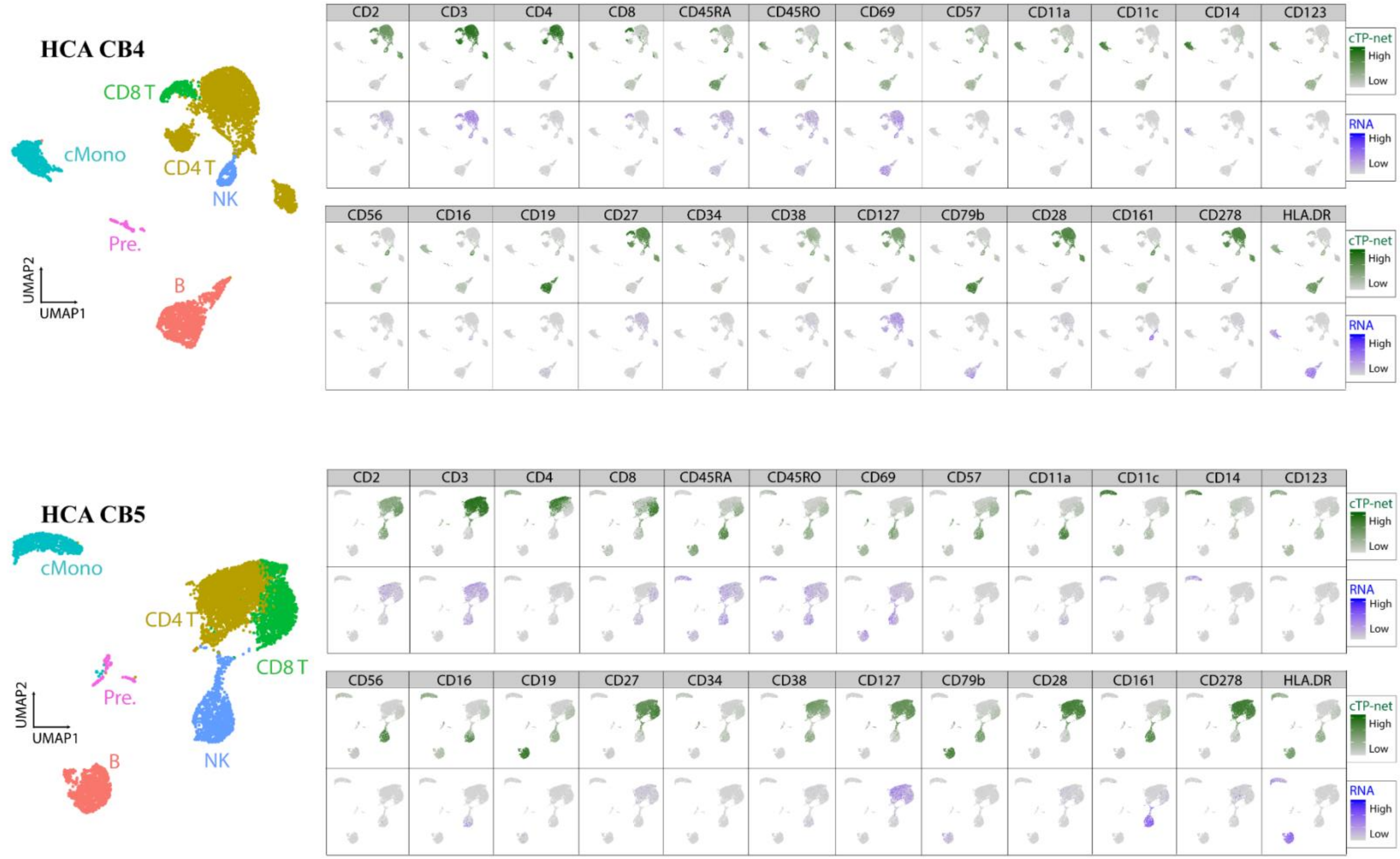

HCA CB4
CD8 T
cMono
CD4 T
NK
Pre.
B
UMAP2
UMAP1
CD2
CD3
CD4
CD8
CD45RA
CD45RO
CD69
CD57
CD11a
CD11c
CD14
CD123
cTP-net
High
Low
RNA
High
Low
CD56
CD16
CD19
CD27
CD34
CD38
CD127
CD79b
CD28
CD161
CD278
HLA.DR
HCA CB5
cMono
CD4 T
CD8 T
Pre.
NK
B
UMAP2
UMAP1

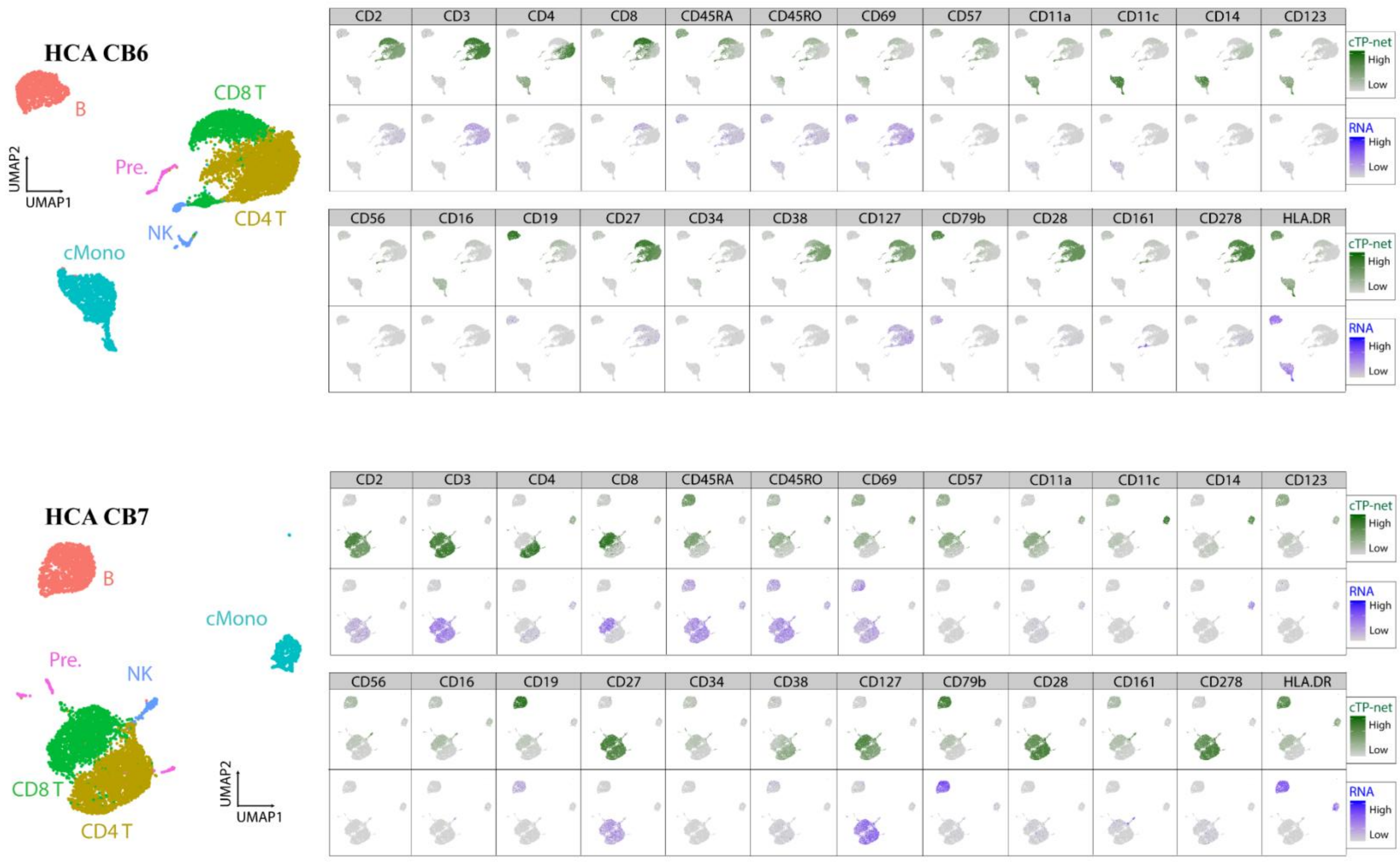

HCA CB6
B
Pre.
CD8 T
CD4 T
NK
cMono
UMAP2
UMAP1
CD2
CD3
CD4
CD8
CD45RA
CD45RO
CD69
CD57
CD11a
CD11c
CD14
CD123
CD56
CD16
CD19
CD27
CD34
CD38
CD127
CD79b
CD28
CD161
CD278
HLA.DR
cTP-net
High
Low
RNA
High
Low
HCA CB7
B
cMono
Pre.
NK
CD8 T
CD4 T
UMAP2
UMAP1

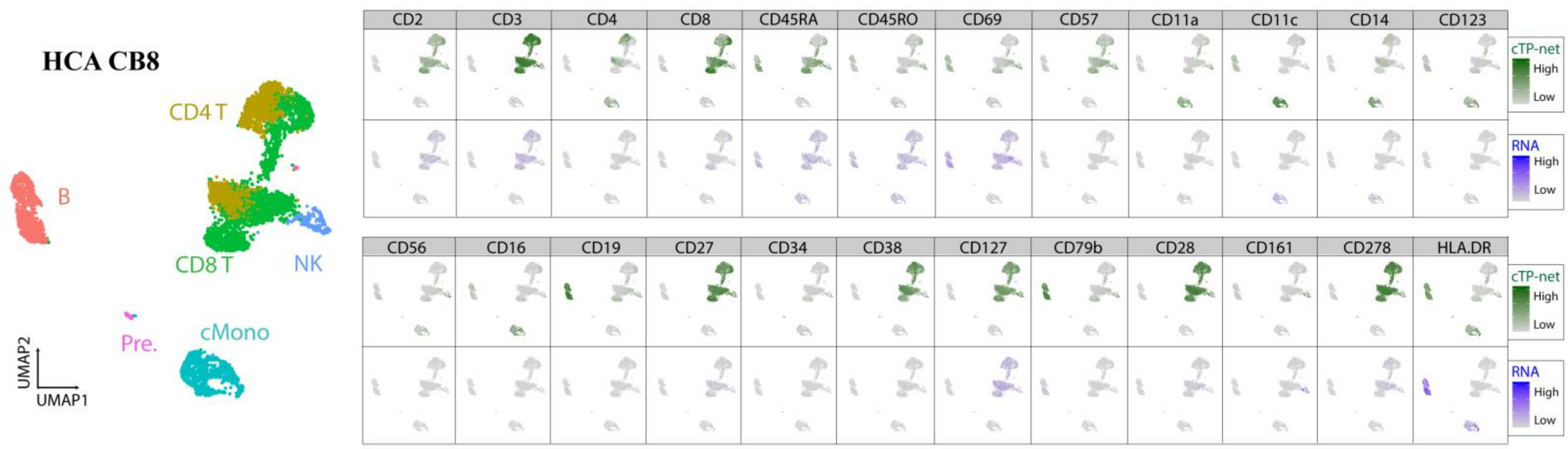
HCA CB8
CD4 T
B
CD8 T
NK
Pre.
cMono
UMAP2
UMAP1
CD2
CD3
CD4
CD8
CD45RA
CD45RO
CD69
CD57
CD11a
CD11c
CD14
CD123
cTP-net
High
Low
RNA
High
Low
CD56
CD16
CD19
CD27
CD34
CD38
CD127
CD79b
CD28
CD161
CD278
HLA.DR
cTP-net
High
Low
RNA
High
Low

| |
|---|
| Supplementary Figure 8 |
| cTP-net prediction on Human Cell Atlas CBMCs by individual. |
| For each individual, we show (1) t-SNE visualization of HCA CBMCs based on expression. B: B cells; CD4 T: CD4 T cells; CD8 T: CD8 T cells; cMono: classic Monocyte; NK: Natural killer cells; Pre.: Precursors. (2) cTP-net imputed protein abundance and RNA of its cognate gene across 24 different surface proteins. |

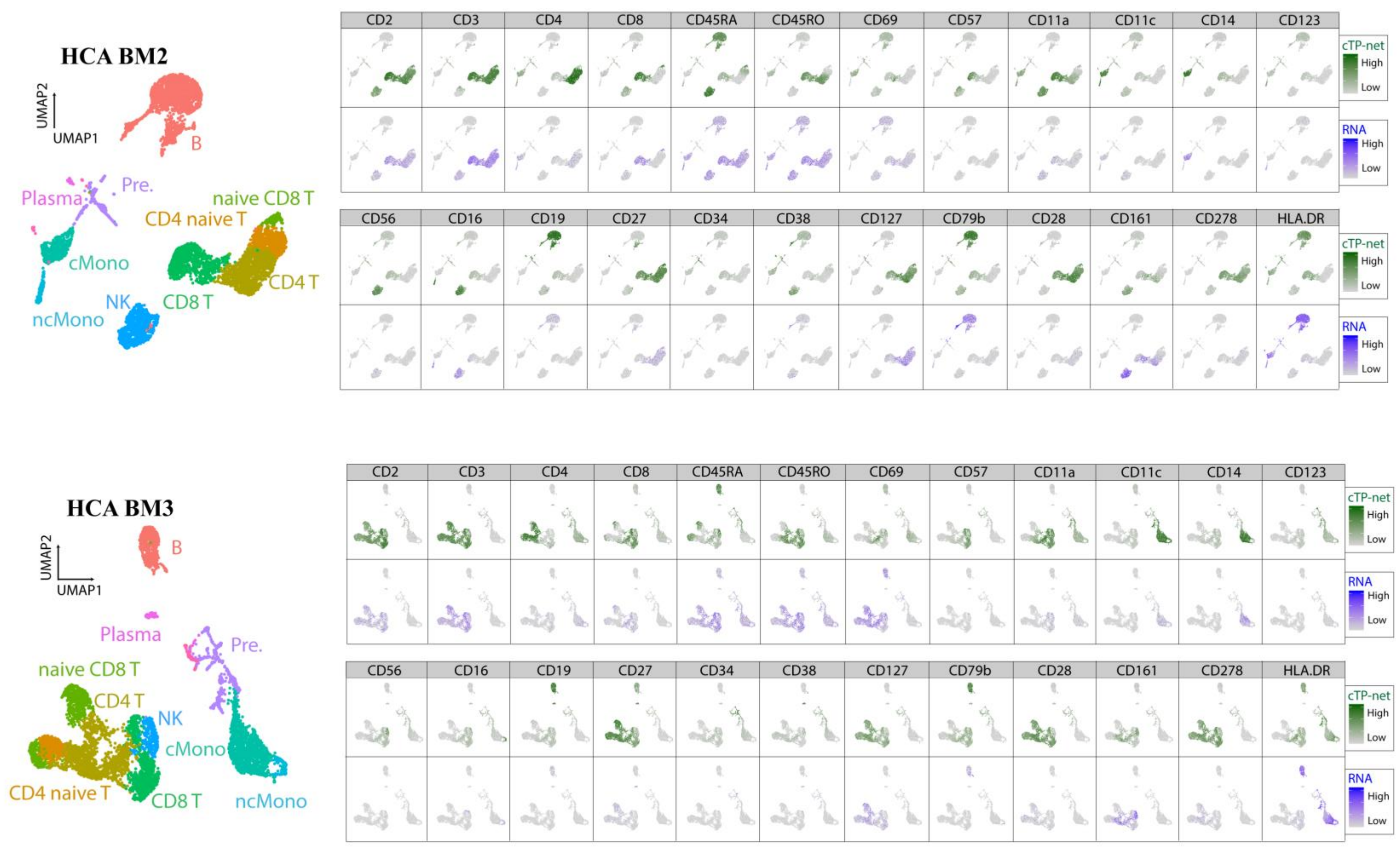
HCA BM2
UMAP2
UMAP1
B
Plasma
Pre.
naive CD8 T
CD4 naive T
cMono
CD4 T
NK
CD8 T
ncMono
CD2
CD3
CD4
CD8
CD45RA
CD45RO
CD69
CD57
CD11a
CD11c
CD14
CD123
cTP-net
High
Low
RNA
High
Low
CD56
CD16
CD19
CD27
CD34
CD38
CD127
CD79b
CD28
CD161
CD278
HLA.DR
HCA BM3
UMAP2
UMAP1
B
Plasma
Pre.
naive CD8 T
CD4 T
NK
cMono
CD4 naive T
CD8 T
ncMono

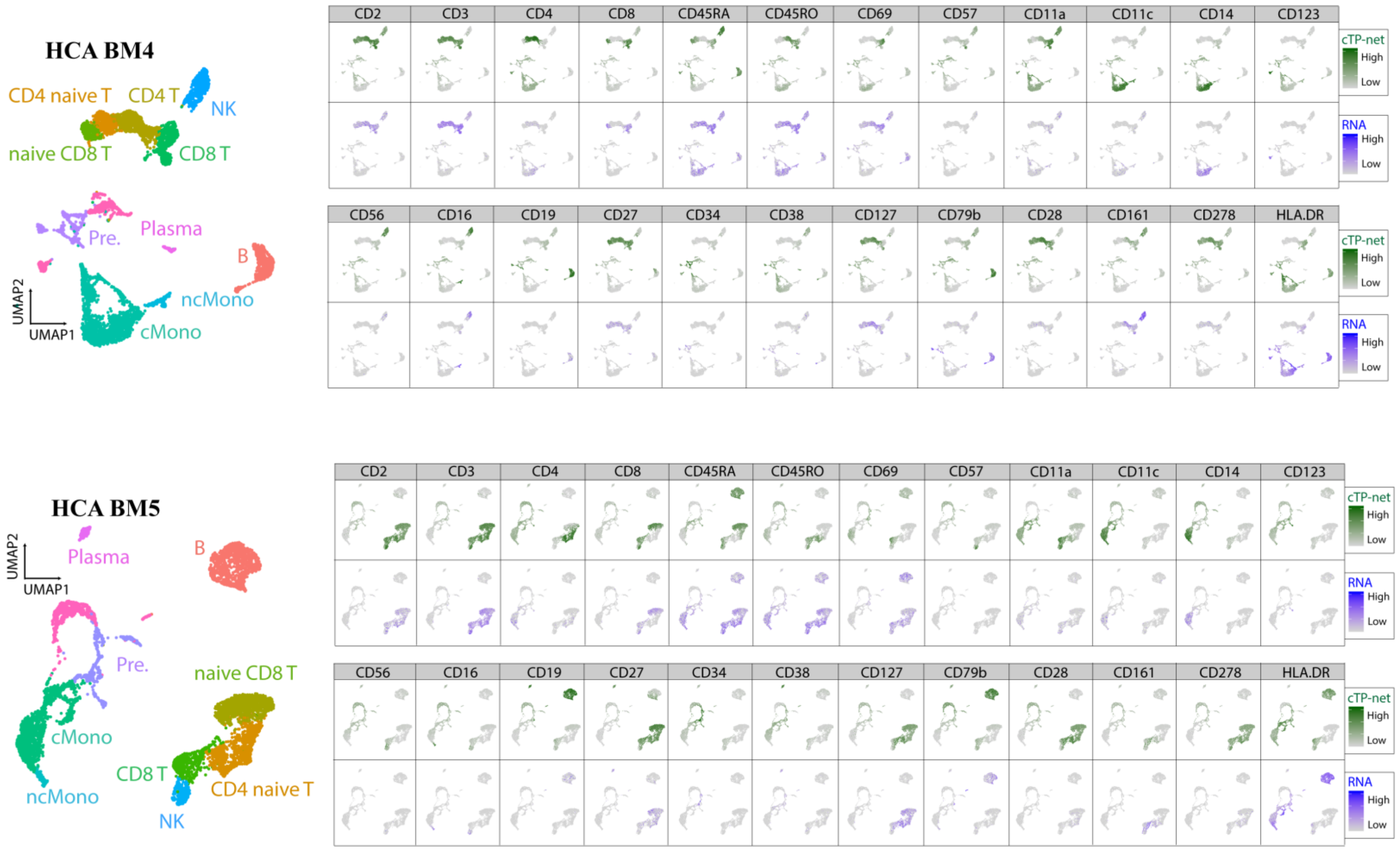

HCA BM4
CD4 naive T
CD4 T
NK
naive CD8 T
CD8 T
Pre.
Plasma
B
ncMono
cMono
UMAP2
UMAP1
CD2
CD3
CD4
CD8
CD45RA
CD45RO
CD69
CD57
CD11a
CD11c
CD14
CD123
CD56
CD16
CD19
CD27
CD34
CD38
CD127
CD79b
CD28
CD161
CD278
HLA.DR
cTP-net
High
Low
RNA
High
Low
HCA BM5
UMAP2
UMAP1
Plasma
B
Pre.
naive CD8 T
cMono
CD8 T
CD4 naive T
ncMono
NK

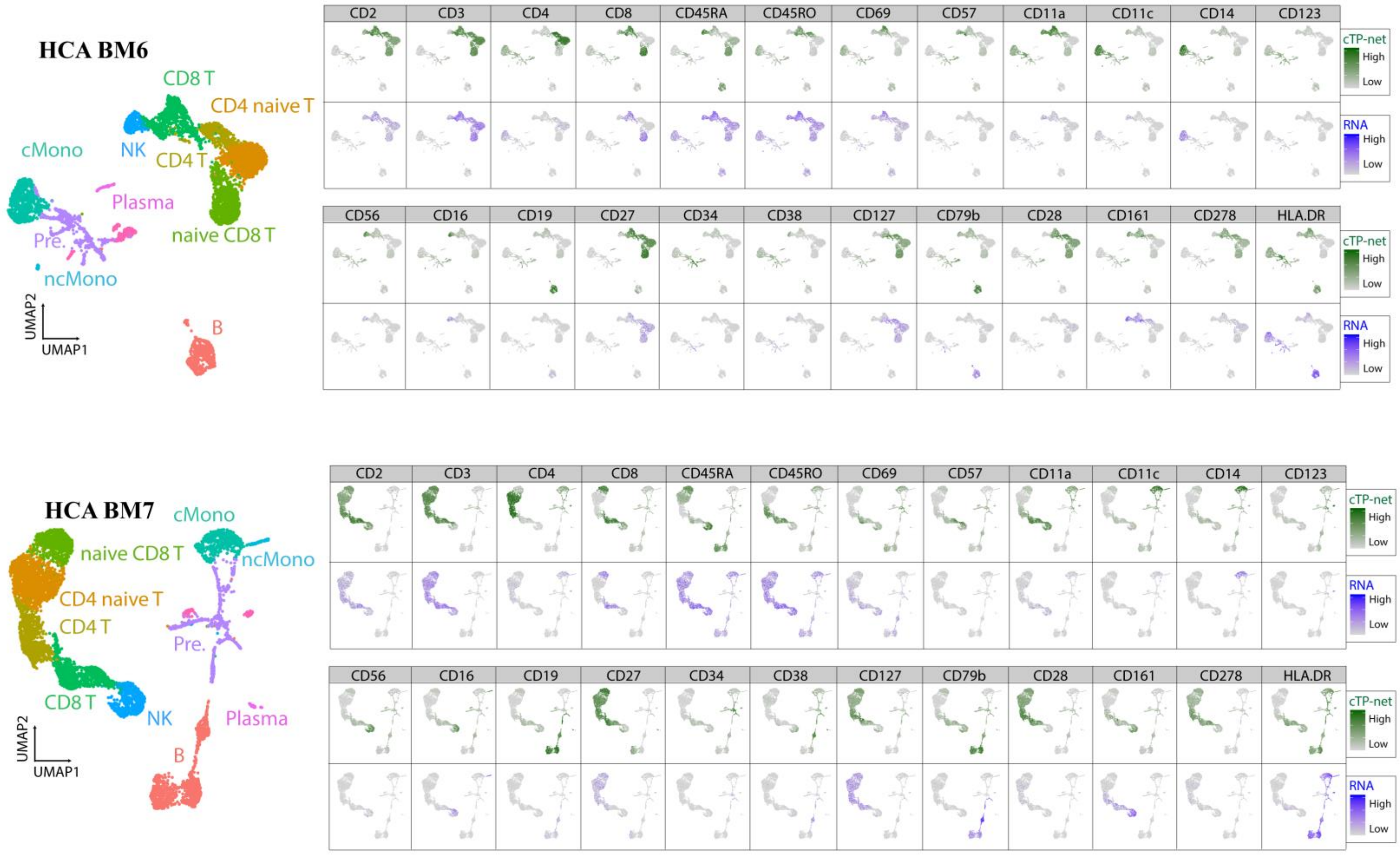

HCA BM6
CD8 T
CD4 naive T
NK
CD4 T
cMono
Plasma
naive CD8 T
Pre.
ncMono
B
UMAP2
UMAP1
CD2
CD3
CD4
CD8
CD45RA
CD45RO
CD69
CD57
CD11a
CD11c
CD14
CD123
CD56
CD16
CD19
CD27
CD34
CD38
CD127
CD79b
CD28
CD161
CD278
HLA.DR
cTP-net
High
Low
RNA
High
Low
HCA BM7
cMono
naive CD8 T
ncMono
CD4 naive T
CD4 T
Pre.
CD8 T
NK
Plasma
B
UMAP2
UMAP1

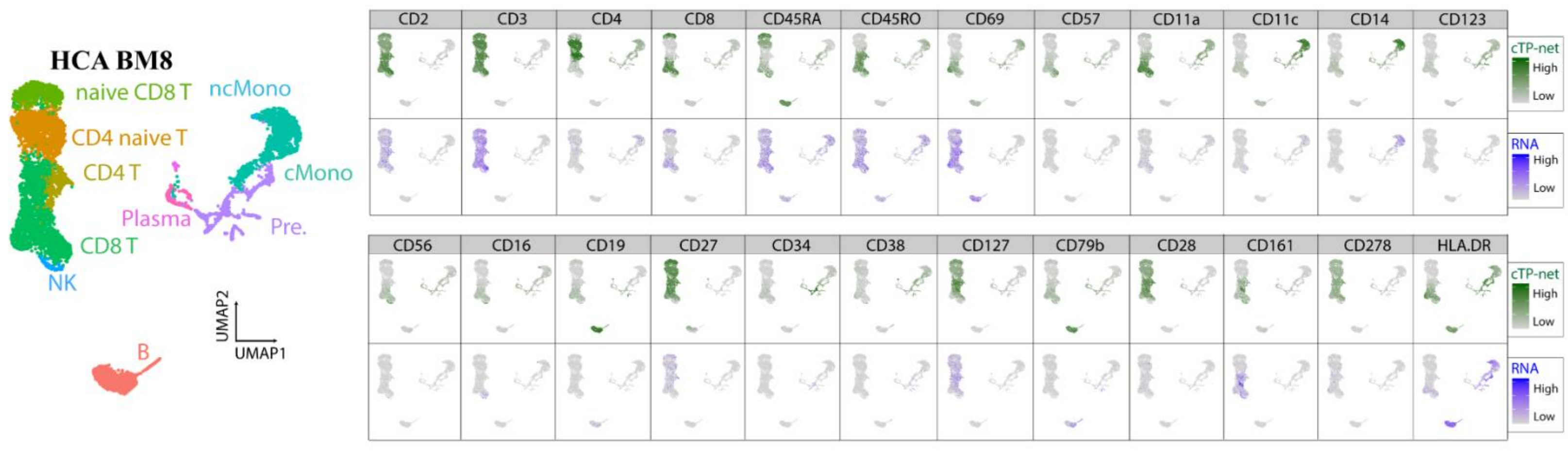

HCA BM8
naive CD8 T
CD4 naive T
CD4 T
Plasma
CD8 T
NK
ncMono
cMono
Pre.
B
UMAP2
UMAP1
CD2
CD3
CD4
CD8
CD45RA
CD45RO
CD69
CD57
CD11a
CD11c
CD14
CD123
CD56
CD16
CD19
CD27
CD34
CD38
CD127
CD79b
CD28
CD161
CD278
HLA.DR
cTP-net
High
Low
RNA
High
Low

| |
|---|
| Supplementary Figure 9 |
| cTP-net prediction on Human Cell Atlas BMMCs by individual. |
| For each individual, we show (1) t-SNE visualization of HCA BMMCs based on expression. B: B cells; CD4 T: CD4 T cells; CD8 T: CD8 T cells; Mono: Monocyte; NK: Natural killer cells; Pre.: Precursors. (2) cTP-net imputed protein abundance and RNA of its cognate gene across 12 different surface proteins. |

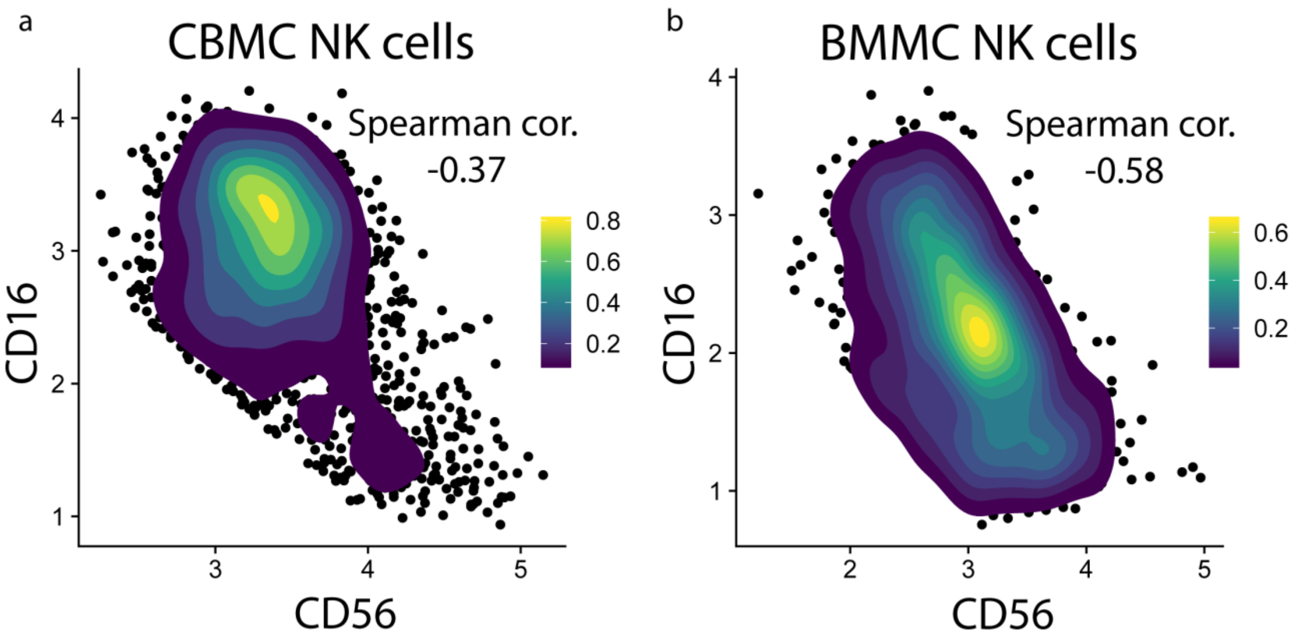


Supplementary Figure 10

Contour plot of cells based on imputed CD56 and CD16 abundance in NK cell populations.

(a) NK cells across all samples from HCA CBMC. (b) NK cells across all samples from HCA BMMC. Strong negative correlation with two subpopulation observed.

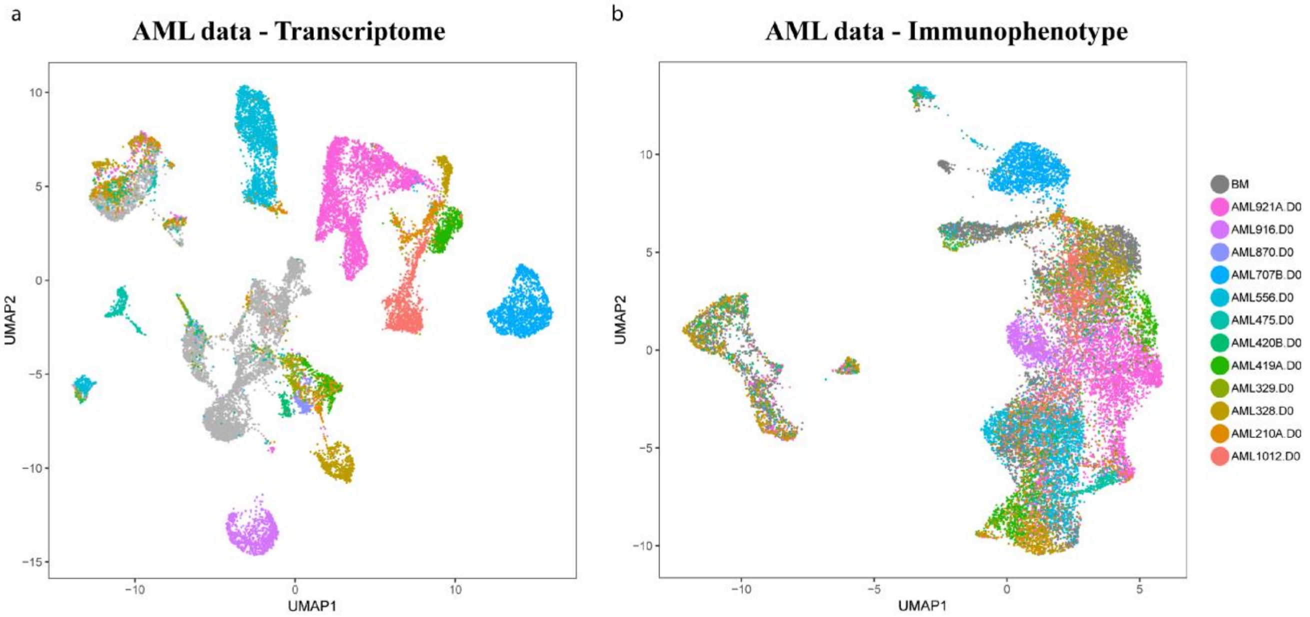


Supplementary Figure 11

UMAP plots of AML data set, colored by samples.

(a) Dimension reduction on transcriptome (RNAs). (b) Dimension reduction on imputed surface proteins.
